# ILEX PARAGUARIENSIS, EXERCISE AND CARDIOPROTECTION: A retrospective analysis

**DOI:** 10.1101/452946

**Authors:** Fábio Cahuê, José Hamilton Matheus Nascimento, Luciane Barcellos, Veronica P. Salerno

## Abstract

Studies on strategies to generate cardioprotective effects have been on the rise. Previous work by our group with an *ex vivo* model of ischemia/reperfusion has shown that both the short-term consumption of yerba mate and exercise can each induce protection of cardiac function independently. Surprising, the two strategies together do not, with an apparent loss of their respective cardioprotection activity. To improve our understanding of the mechanisms involved without reperforming the experiments, we have conducted a retrospective data science-analysis that have produced new insights. The analysis shows that yerba mate generated reductive stress. Alone, this stress increased redox damage in the heart that appears to have led to a protective conditioning. In combination with exercise, the effects of mate inhibited the intermittent ROS generation promoted by exercise alone, which diminished the adaptive response in the heart. These results suggest that an understanding of the molecular mechanisms involved with the yerba mate-promoted reductive stress in cardiac tissue could lead to improved strategies to induce cardioprotection.

## Introduction

Cardioprotective strategies are an important approach to prevent morbidity and death from myocardial infarctions (heart attacks). This is reflected in the increased number of studies on methods to provide protection against ischemia/reperfusion injuries in the heart. A recent search of PubMed, performed 09/05/2018, with “cardioprotection” as the keyword returned 3801 articles since 01/01/2010. This represents a 49% increase over the previous, equivalent period (2536 articles from 2000 to 2009).

Our recent work described the cardioprotective effect of short-term consumption of *Ilex paraguariensis* (St. Hill) (Cahue et al. 2017). The results showed that the daily ingestion of *I. paraguariensis* (1 g/kg) for seven days could promote a protection of cardiac function against a global ischemia/reperfusion (I/R) injury. The measured increases in protein carbonyls and lipid peroxidation in the cardiac tissue of animals provided *I. paraguariensis* suggested a possible oxidative preconditioning mechanism. However, the combination of *I. paraguariensis* consumption with low-intensity aerobic exercise, which is cardioprotective, attenuated the benefits of each. This counteractive effect was proposed to be related to the antioxidant effect of *I. paraguariensis* attenuating the oxidative burst associated with exercise, which inhibited the normally observed increase in superoxide dismutase (SOD) activity that is promoted by exercise.

A slight, but significant increase in the GSH/GSSG ratio in cardiac tissue of rats that consumed *I. paraguariensis* suggested that a pro-oxidative preconditioning could be promoted by its short-term consumption. Here, we have reanalyzed the original data set (Cahuê et al., 2017) using data science (DS) methods, which some authors describe as the science that learns from data (Donoho 2017). The results help clarify possible mechanisms underlying *I. paraguariensis* and exercise induced cardioprotection and the inhibition caused by the interaction between the herb and exercise

## Interpreting Graphical Analysis

Graphics were generated through an online tool, RAWGraphs (http://app.rawgraphs.io/), that can utilize raw data. The data from our previous work (Cahuê et al, 2017) was extracted, organized and saved in an ․csv file to be compatible with RAWGraphs. To preserve the balance in the analyses, which can be influenced by the number of samples per group, a sample size of five was used per group for all. The variables used in all graphical analyses were TBARS (lipid peroxidation), PC (protein carbonyls), SOD (Superoxide Dismutase activity) and GSH/GSSG ratio based on their significance to the results of our previous work.

Three types of graphic representations were built to analyze the data graphically. Parallel Coordinates plots provided an equalized scaled, multivariable graphic to allow the visualization of patterns in the data. Alluvial Diagrams shows the contribution level of a variable within each group, attributing a “weight” based on a main variable. This graphic displays lines that are ordered from higher (thicker) to lower (thinner) values converging on an impact of values in each group. Finally, a Convex Hull shows, in a dispersion graphic, a polygon-based grouped analysis that helps to understand the behavior of each group when correlating two variables. All three graphics, together, can show patterns that can be used to predict which variables are related with the correspondent outcome (in our case, cardioprotection).

Figure 1 shows a Parallel Coordinates graphic using redox damage biomarkers, lipid peroxidation (TBARS) and protein carbonylation level, a redox balance biomarker, reduced/oxidized glutathione ratio (GSH/GSSG), and superoxide dismutase (SOD) activity. We focused our analysis on the mate consumption only group (M; black line) and the exercise training only group (E; red line). An examination of the characteristics of the data lines shows that the samples of the M group displayed higher values of PC, TBARS and GSH/GSSG ratio without a pattern in SOD activity. In samples from the E group, values of SOD activity were higher than those observed in M group, with lower values for PC and TBARS. These observations suggest that *I. paraguariensis*-induced cardioprotection, in our model, must be related with higher redox damage and a slightly higher redox balance.

**Figure 1.**
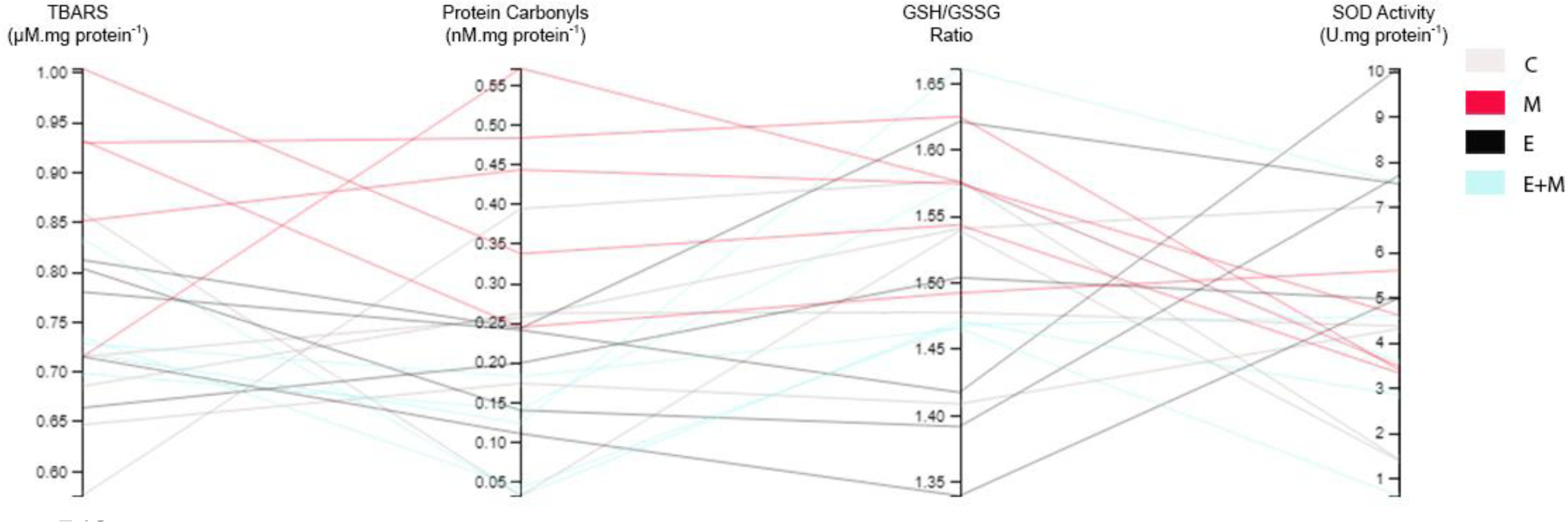
Parallel Coordinates composed by TBARS, PC, GSH/GSSG Ratio and SOD activity. Black lines – Mate group. Red lines – Exercise Group. Light Blue lines – Exercise + Mate group. Light Grey lines – Control group. For each group n=5.

Panel 2 shows how much impact these variables exerted in each group. Both redox damage biomarkers (PC in figure 2A and TBARS in figure 2B) and GSH/GSSG ratio (figure 2C) had a higher impact in the M group. TBARS had a moderate impact in the E and E+M group, while PC showed a lower impact in E and E+M. In contrast, SOD activity can be observed as a main effect of exercise intervention (figure 2D).

**Figure 2.**
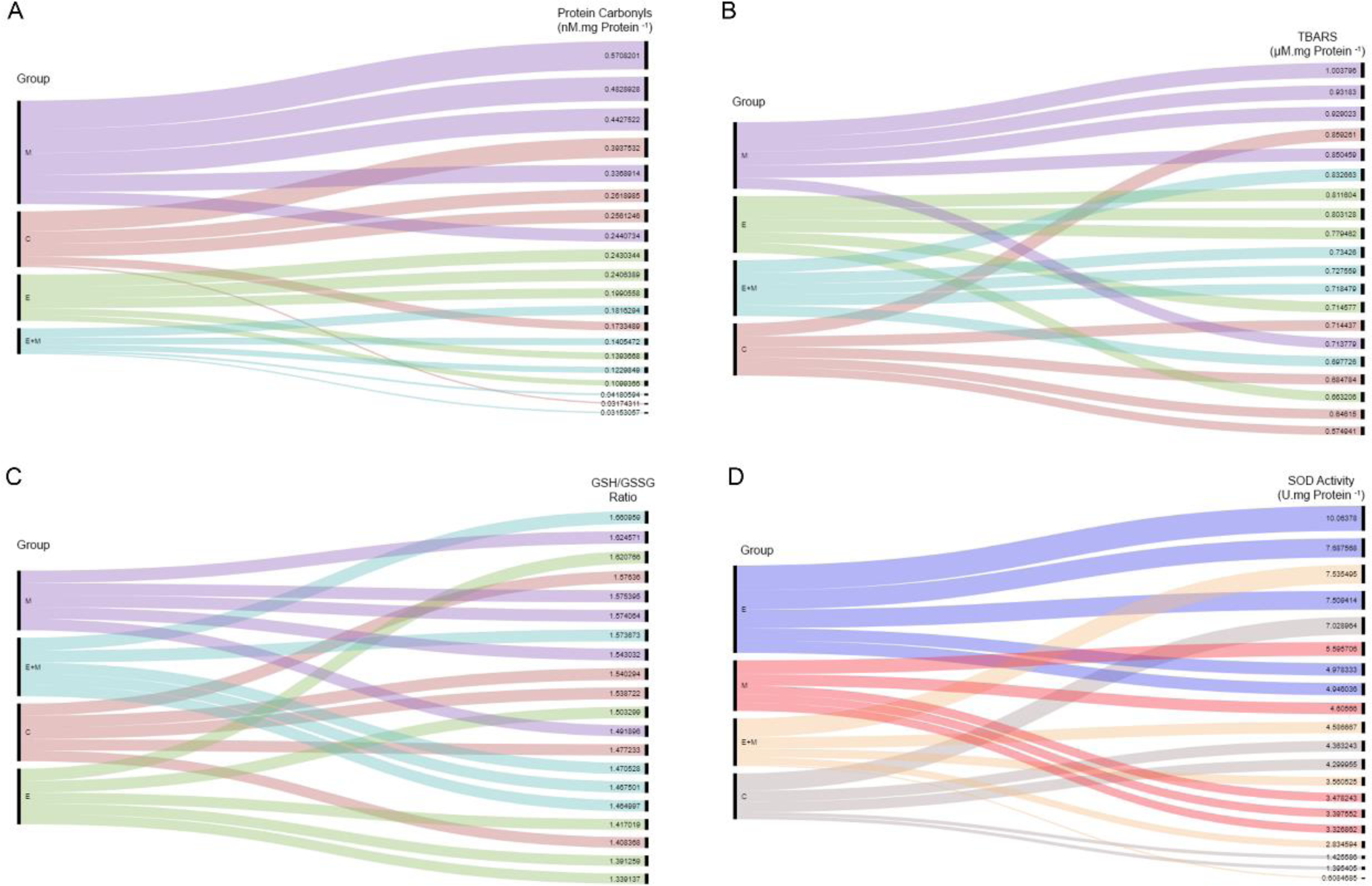
Alluvial Diagram showing the impact of PC (A), TBARS (B), GSH/GSSG Ratio (C) and SOD activity (D) in each group. The order of group, from top to bottom, reflects the impact of individual values in the respective group. For all graphics, n=5 for each group.

In figure 3A and 3B, there can be observed within the M group a pattern with the samples being positioned in the up-right corner representing higher levels of PC (3A), TBARS(3B) and GSH/GSSG ratio, which is consistent with the behavior observed previously (Figures 1, 2A and 2B). To verify the impact of SOD activity in PC and TBARS, we correlated these variables and E group samples showed higher values of SOD activity, accompanied with lower values of PC (figure 3C) and TBARS (3D). Group E+M showed lower and intermediate values of SOD activity with lower levels of PC and TBARS, suggesting the antioxidant and inhibitory effect of *I. paraguariensis* consumption on the adaptations from exercise training.

**Figure 3.**
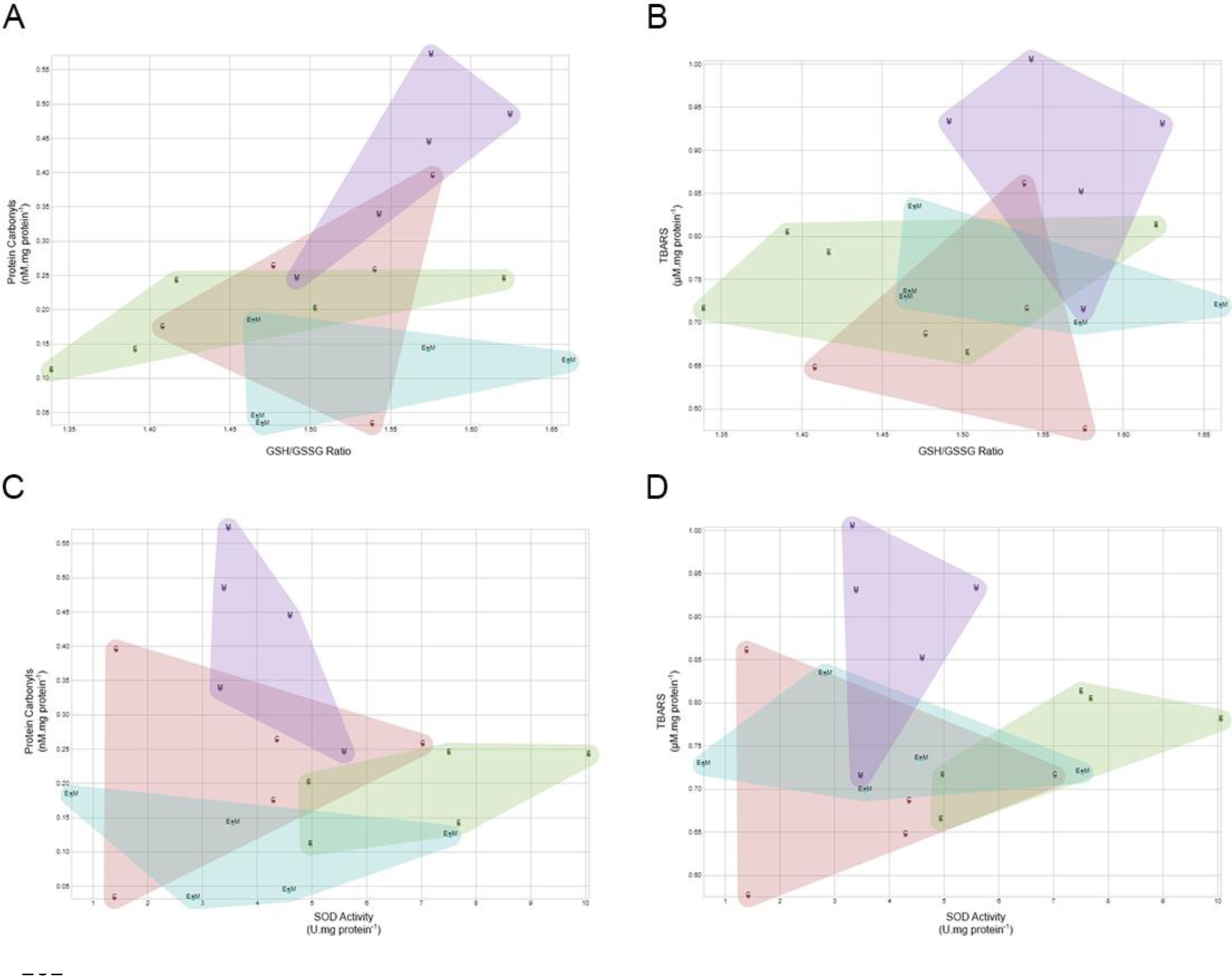
Convex Hull graphics, correlating PC and TBARS with GSH/GSSG Ratio (3A and 3B), and with SOD activity (3C and 3D). For all graphics, n=5 for each group.

## A New Approach for *Ilex paraguariensis*-mediated Cardioprotection

In our previous work, the mechanism we proposed for the cardioprotection qualities of *I. paraguariensis* consumption was through the pro-oxidative effects expected at the concentration provided (Miranda et al. 2008), which could induce a preconditioning adaptation. Another possible explanation to explain the effects observed in our study is reductive stress (RS). The phenomenon of RS was initially associated with the effects from the large amount of antioxidant, either endogenous or exogenous, which modulated signaling pathways that diminished the antioxidant capacity or mitochondrial function (Passi et al. 1987). Recently, RS has been described to coincide with an increase in the reductive capacity due to a huge increase in ratios of GSH to GSSG, NADH^+^H^+^ to NAD or NADPH^+^ to NADP ratios (Perez-Torres et al. 2017). It also can occur when an antioxidant compound interacts with other molecules that generates free radicals or improves the production of superoxide.

A prolonged exposure to RS can induce hypertension, DNA damage, apoptosis (Hodnick et al. 1986), mitochondrial dysfunction and endothelial cell death (Posadino et al. 2013; Posadino et al. 2015). *I. paraguariensis* contains high concentration of phenolic compounds (Bracesco et al. 2011) that can induce to ROS production when they interact with transition metal ions (i.e. copper and iron) or when they are converted into phenoxyl radicals in a high reductive environment (Passi, Picardo et al. 1987; Decker 1997). This effect can be related with others already explained in literature. Du et al. (2017) showed an increase in peroxide production following an increase in NOX4 and p22^Phox^ protein expression in rat mesenteric venules after a single injection of 7 mg/kg chlorogenic acid, a phenolic acid and a major constituent of *I. paraguariensis* (Bracesco, Sanchez et al. 2011). This dose corresponds to a 5-fold increase in the dose indicated by Chinese herbal medicine injection protocols. Murakami et al. (2013) quantified the concentration of phenolic acid in 30 mg/mL of dried leaves of *I. paraguariensis* and showed a concentration of about 1.99 mg/mL of chlorogenic acid. de Oliveira et al. (2017) showed an increase of dyhydrophenylproprionic acid and hippuric acid (phenolic acids derived from chlorogenic acid metabolism) concentration in liver and plasma 8 hours after a single treatment with 2 g/kg of *I. paraguariensis*. In our experimental model we used a dose of 1 g/kg of a commercially preparedlyophilized extract of *I. paraguariensis* (Matte Leão, Curitiba, Brasil). Each animal consumed about 200~300 mg/mL of this extract which represent a phenolic acid concentration between 13~20 mg/mL, which correspond to 9-14-fold the dose indicated in Chinese herbal medicine injection protocols. Thus, it is expected that this higher concentration of yerba mate, for 7 days, could potentiate the reductive environment, inducing to the reductive stress.

The potential of short-term *I. paraguariensis* consumption to induce RS can explain the observed inhibition of cardioprotection when combined with low-intensity aerobic exercise. Several studies have shown similar inhibitory effects from the combination of exercise training and antioxidants supplementation. Gomez-Cabrera et al. (2008) found a negative interaction of vitamin C (500 mg/kg bw/day) and exercise training (5 days per week, 25-25 minutes per day) for 3 weeks in running capacity and Cytochrome C expression. A combination of 12-week endurance training with vitamins C and E supplementation in humans increased plasma protein carbonyls (Yfanti et al. 2012), which represents a negative effect from their interactions. Our data showed an inhibition of SOD activity in the exercise training group that also consumed *I. paraguariensis*, which resulted in an inhibition of recovery during the reperfusion period. Therefore, our results converge with previous observations in the literature that were exposed by the data science approach. It suggests that there is a negative interaction between antioxidant supplementation and exercise training that impacts myocardial tissue. Further studies are warranted to better understand this phenomenon.

## Conclusion and a New Perspective

The retrospective analysis of data by a data science approach suggests that *I. paraguariensis*-mediated cardioprotection can be explained by a reductive stress. This condition can be responsible for the increased levels of lipid peroxidation and protein carbonyls as well as the inhibition of exercise-mediated cardioprotection. This effect can be mainly attributed to the high concentration of polyphenols contained in yerba mate consumed during the experimental period. Our group continues to concentrate our efforts to understand the molecular mechanisms underlying these effects with a focus on the interaction between the redox damage and the regulation of mRNA expression of genes related to antioxidants and the activity of antioxidant enzymes to clarify how *I. paraguariensis* promotes cardioprotection and inhibits exercise-related protection against I-R injury.

## Acknowledgments

Dr. D. William Provence Jr. gave a careful attention to English during revision of the manuscript. This work received financial support from Coordenação de Aperfeiçoamento de Pessoal de Ensino Superior (CAPES), Fundação de Amparo à Pesquisa do Estado do Rio de Janeiro (FAPERJ), and Conselho Nacional de Desenvolvimento Científico e Tecnológico (CNPq).

